# Perceiving Material Qualities from Moving Contours

**DOI:** 10.1101/2025.07.21.665763

**Authors:** Amna Malik, Ying Yu, Huseyin Boyaci, Katja Doerschner

## Abstract

While research on the perception of line drawings has long demonstrated the importance of contours in object recognition, recent work shows that contours can also convey material properties. For example, even simple 2D shapes with varying contours have been shown to evoke vivid impressions of different materials (Pinna & Deiana, 2015). However, such static representations capture only a single moment in time. When a material moves, its contours shift, evolve, or deform over time, creating contour motion. Does this contour motion convey diagnostic information about material properties, independent of surface appearance? Existing studies on the role of dynamic cues in material perception either use fully rendered 3D stimuli, where contour motion is confounded with rich surface information, or motion-only displays (dynamic dot stimuli or noise patches), which eliminate surface cues but also lack clearly defined contours. As a result, the relative contribution of contour motion to material perception remains unclear. To address this gap, we measured how human observers perceive materials from dynamic line drawings (“line”), compared to animations of fully textured stimuli that carry optical and motion information (“full”), as well as dynamic dot stimuli (“dot”). Stimuli were rendered versions (full, dot, line) of material animations from five material categories (jelly, liquid, smoke, fabric, and rigid-breakable). In one experiment, participants rated five material attributes (dense, flexible, wobbly, fluid, airy motion), and in a second experiment, participants were asked to choose one of the two materials that is more similar to a third material across all possible combinations. Results from both experiments consistently reveal that 1) Dynamic line drawings vividly convey mechanical material properties, and 2) the similarity in material judgments between line and full conditions was larger than that between dot and full conditions. We conclude that contour motion carries rich information about mechanical material qualities.

## Introduction

Lines play a fundamental role in visual perception. This is evident from the earliest known cave drawings, dating back at least 45,500 years, where early humans used lines to depict objects, animals, and scenes from life (Oktaviana et al., 2024). The persistence of line-based representation across human history suggests that our visual system is attuned to processing lines as an essential cue for recognizing shapes, forms, and structures in our environment. Research shows that humans can recognize objects and scenes from line drawings with remarkable efficiency, even in the absence of shading, texture, or color cues (Biederman & Ju, 1988; Elder, 2018, 2018; Elder & Goldberg, 2002; Greene & Oliva, 2009; Marr & Nishihara, 1978; Sann & Streri, 2007). The question is, what makes line drawings so effective? To answer this, we should consider what lines represent. Line drawings primarily correspond to meaningful edges (contours) in real-world images. In the absence of surface texture, contours of an object can correspond to: occluding contours (such as smooth occlusions, where the surface of the object curves away from the viewer, crease occlusions, where surfaces meet at a sharp edge), surface creases (folds or ridges that indicate internal shape structure), or illumination-related contours (like shadows and specular highlights). While all of these can be depicted in line drawings, studies have shown that occluding contours and creases are especially informative for recognizing object shape and identity (Bansal et al., 2013; Koenderink, 1984).

Since object recognition from line drawings is very robust, it has inspired edge-based theories of object recognition, such as those proposed by Biederman (1987). According to these accounts, contours provide the critical structural information needed for identification, while surface properties such as texture or color play a minimal role. Material perception, in contrast, has placed comparatively less emphasis on the role of contours as surface cues appear to be central for estimating and recognizing material properties (Fleming, 2014). With few exceptions, the role of contours in material perception has received little attention. Pinna and Deiana (2015) provided compelling evidence that contours alone provide substantial information about material properties. In one of the experiments, they presented 2D squares depicted with various contour shapes and asked observers to identify the material. Participants spontaneously attributed distinct material identities to each square based on the shape of the contour alone, e.g., straight edges were seen as plastic, wavy contours as fabric, and sharp spikes as metal. In another experiment, they showed multiple squares to participants with each square retaining a clean outline except for one locally distorted edge (Pinna & Deiana, 2015). These subtle distortions, jagged notches, rounded bulges, or irregular outflows, led observers to infer events like tearing, melting, or spilling. Remarkably, these static line drawings elicit vivid impressions of what the object is made of and what has happened to it based on contour variation (Schmidt et al., 2016; Spröte et al., 2016; Spröte & Fleming, 2013).

However, it’s important to consider that real-world materials are seldom static; instead, they frequently change as they interact with external forces. While some, such as glass or brittle objects, break upon impact, others, like rubber or jelly, temporarily deform before returning to their original shape when interacted with. Moreover, inherently dynamic materials like liquids and smoke continuously change shape, even in the absence of direct external forces. As a result, contours evolve over time, reflecting characteristic changes unique to each material, and these changes may provide an even richer source of information, capturing aspects of material behavior that static line drawings cannot convey. For example, consider Figure 1(left panel), which shows one of the frames extracted from an animation. This image can be interpreted in multiple ways: it might resemble a doughnut, a bean bag, a coffee bean, or a tomato. However, when the stimulus is viewed dynamically across multiple frames, the material identity becomes immediately clear as the contours move (right panel of Figure 1), namely a balloon, made of a very soft fabric.

**Figure 1.**
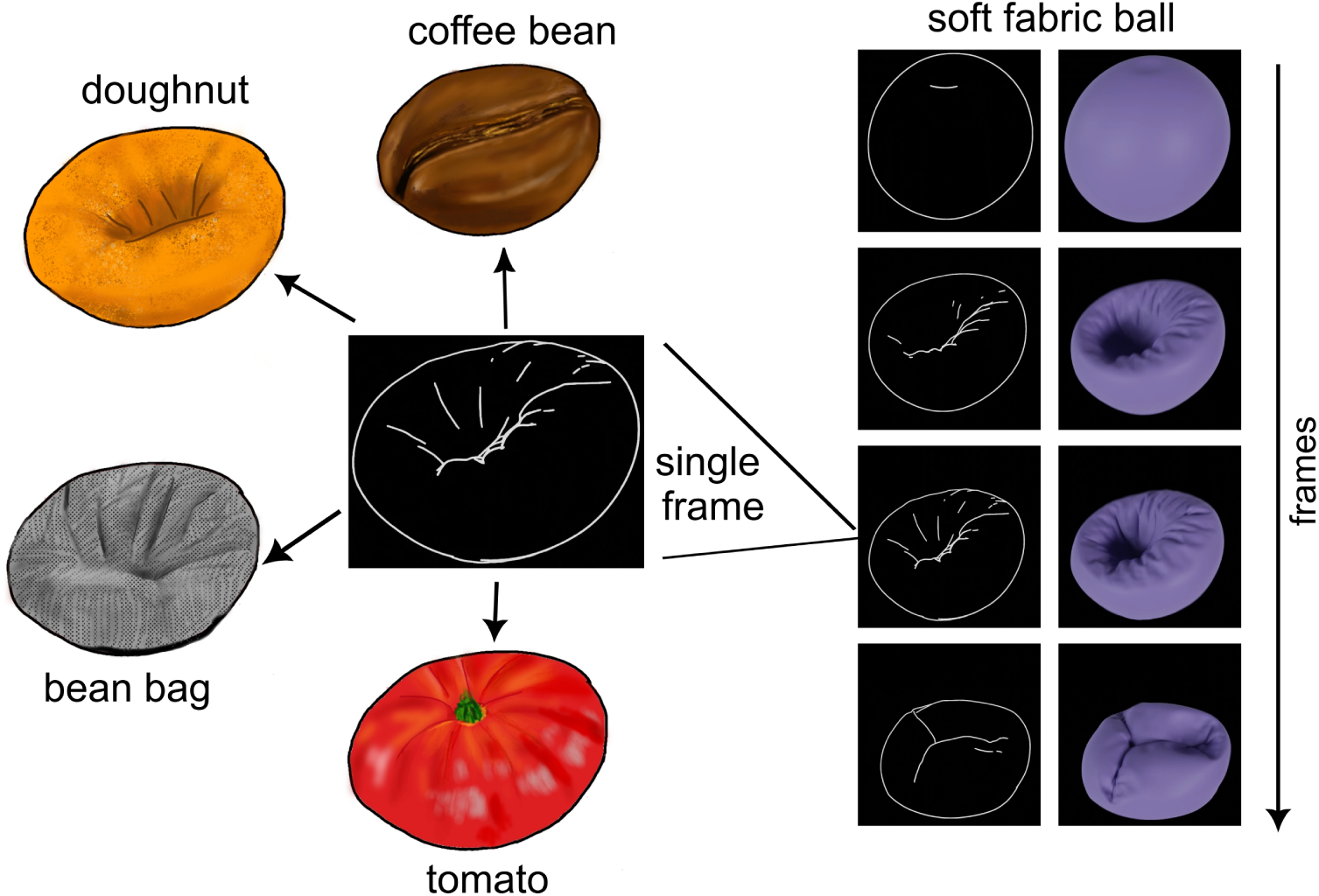
A static frame extracted from an animation depicting a soft fabric can be interpreted ambiguously as different objects. However, when viewed dynamically across multiple frames, contour motion makes the material identity immediately apparent.

Here we test whether two cues emerging from contour motion, shape changes over time, and motion cues such as speed, acceleration, etc., can explain how humans perceive material qualities from line drawings. Previous research shows that both shape and motion cues play a role in the perception of material qualities (Bi et al., 2018, 2019; Kawabe et al., 2015; Kawabe & Nishida, 2016; Paulun et al., 2017; Schmid & Doerschner, 2018; van Assen & Fleming, 2016). However, these studies either use fully rendered 3D stimuli, where contour motion is confounded with rich surface information, or motion-only displays (dynamic dot stimuli or noise patches), which eliminate surface cues but also lack clearly defined contours. As a result, the specific contribution of contour motion to material perception remains unclear. One notable exception is Kawabe & Nishida (2016), who isolated contour motion (only smooth occlusions) by removing specular and diffuse reflections from a jelly cube’s surface. Their findings showed that elasticity judgment was influenced by contour deformation, but their study was limited to a single material (jelly) and a single property (elasticity). We will investigate whether observers are able to reliably infer a broader range of material properties across different materials using only contour motion cues.

To isolate contour motion, we use dynamic line drawings (line drawing animations), rendering only contours (both occluding contours and creases) of an object. We also rendered a dynamic dot version of stimuli as used by Schmid & Doerschner (2018) and Bi et al. (2019). Dynamic dot stimuli depict material motion by dots stuck in or on the surface of the material, thereby removing optical cues. They primarily provide internal motion cues, where each dot changes position over time, generating apparent motion signals through its trajectory. The instantaneous positions of dots at any given moment define the object’s overall structure. Temporal integration of this structural information enables perception of how the object’s 3D shape and surface evolve over time. To some extent, dynamic dot stimuli also convey changes in occluding contour, although much weaker than line-based representations (see Figure 2). In contrast, dynamic line drawings make boundary shape changes explicit by directly representing occluding contour motion (OCM). In addition, they also provide information about how contours evolve over time within the object’s boundary (internal creases), referred to as internal contour motion (ICM). Together, these characteristics of line drawings make them a reliable source of contour motion. Here, we investigate the relative contribution of contour motion in material perception using these dynamic line drawings. Specifically, we ask whether contour motion conveyed by line drawings leads to material perception similar to that evoked by full-textured stimuli, and if so, how does it compare to dot stimuli?

**Figure 2.**
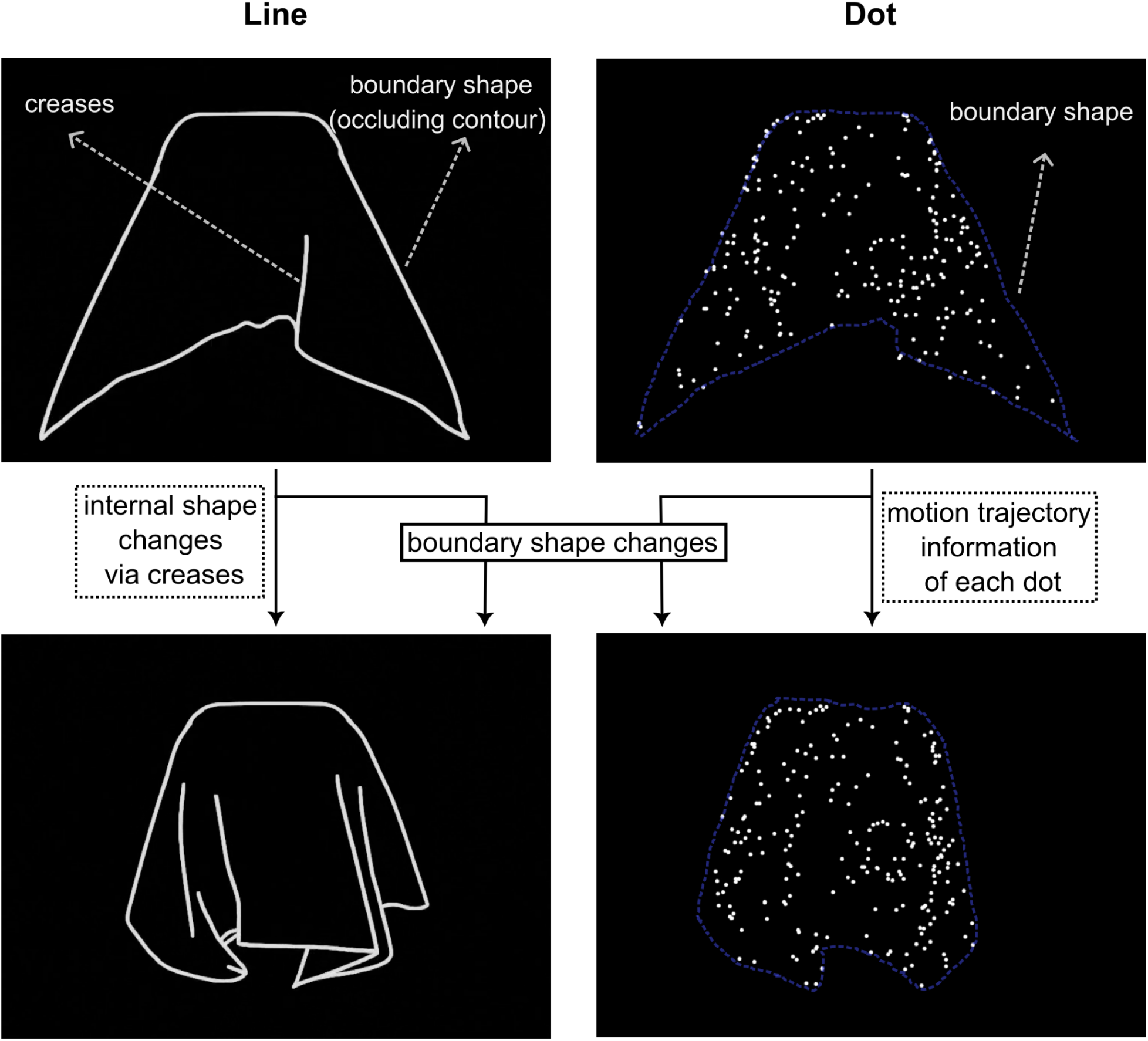
Representative frames from one of the animations are shown for both the line and dot conditions. Line drawings depict occluding contour motion (conveying boundary shape changes) and crease/internal contour motion (showing how contours evolve within the object’s boundary). Dot stimuli convey internal motion cues through the trajectories of individual dots and provide coarse structural information from their instantaneous positions. The blue dotted line in the dot condition is included for illustrative purposes to highlight how boundary shape changes can also be inferred from the dot stimuli, although this cue is weaker than in the line condition.

To test this, we created computer-generated animations representing five material categories: jelly, liquid, smoke, fabric, and rigid-breakable, as shown in Figure 3. These animations featured randomly shaped objects reacting to external forces, exhibiting their dynamic behaviors such as deformation, motion, or dispersion. We rendered each animation in three versions: line, full, and dot. We conducted two experiments: a rating experiment using a semantic task in which participants rated five material attributes (dense, flexible, wobbly, fluid, airy motion), and a similarity matching experiment using a non-semantic task, in which participants viewed animations of materials in a triplet-alternative forced-choice setting and were instructed to choose one of the two materials that was more similar to the third material across all possible combinations. The latter experiment served to validate whether the chosen attributes sufficiently capture the perceptual space. Results from both experiments suggest that contour motion plays a critical role in the perception of mechanical material qualities. In order to explore what information our visual system extracts from these dynamic stimuli, we estimated several motion and shape statistics for line and dynamic dot stimuli and correlated them with the perceptual data. We found that motion or shape-derived statistics can only partially explain perceptual differences across material categories. This suggests that more work is needed to discover and quantify visual features that drive the perception of material qualities in dynamic scenes.

**Figure 3.**
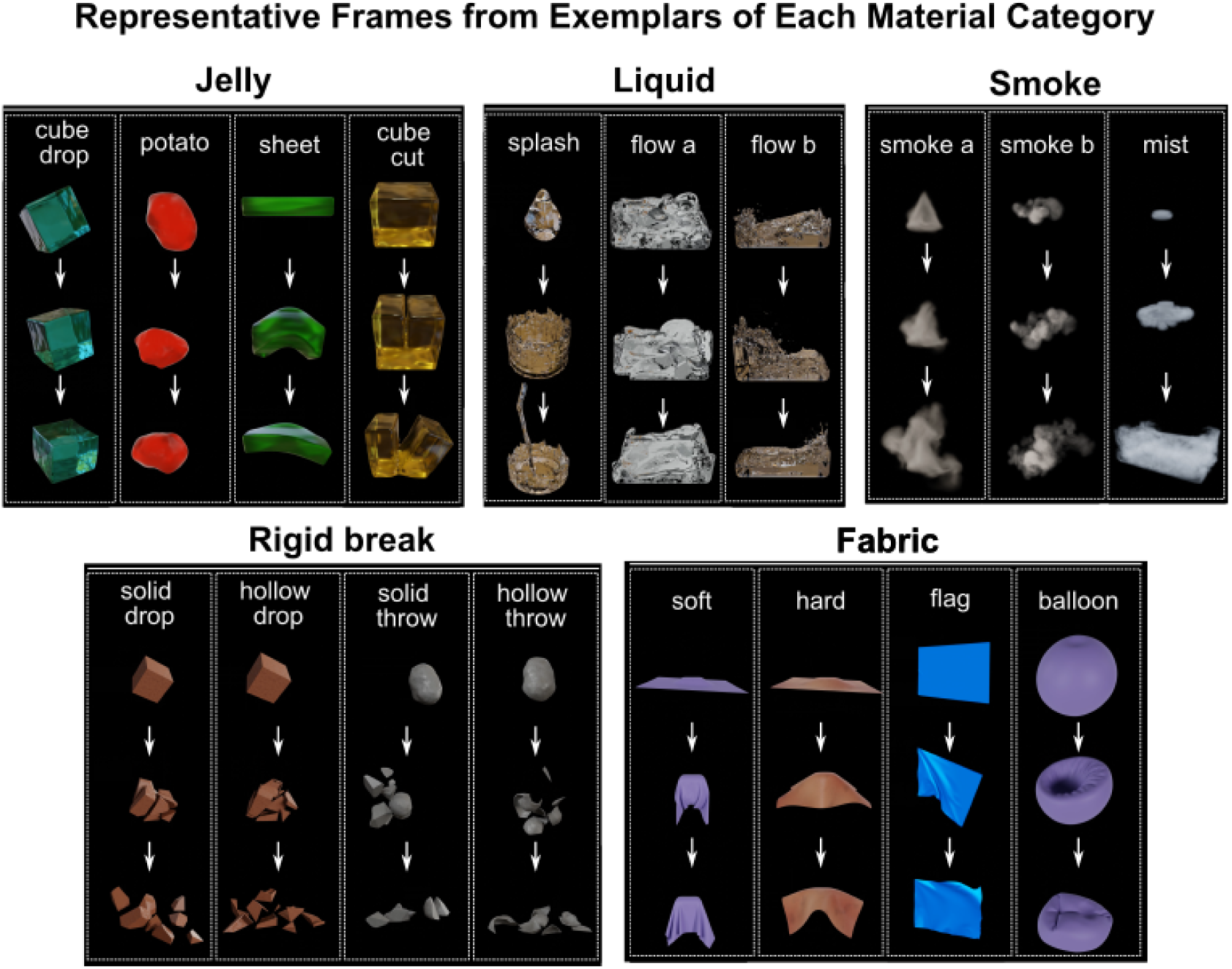
The stimulus set comprises 18 exemplars, each belonging to one of the five material categories: jelly, liquid, smoke, fabric, and rigid-breakable. Three representative frames (frames 3, 15, and 40) from the total of 48 frames for each material category are shown, with arrows indicating their temporal order. While each animation was rendered from 6 to 8 camera angles, only one camera angle is displayed here.

## Materials and Methods

### Stimuli

To create a stimulus set with a sufficiently broad range of material classes, we rendered animations of materials from five distinct categories: jelly, liquid, smoke, fabric, and rigid-breakable. Each category contained multiple animations representing different variations within the same material class, which we refer to as *exemplars*. Specifically, the jelly, fabric, and rigid-breakable categories each included four exemplars, while the liquid and smoke categories each contained three exemplars. Figure 3 shows representative frames (frames 3, 15, and 40) from exemplars of each material category. All animations were rendered from six or eight camera angles, yielding 114 animations in total. Each animation comprised 48 frames, presented at a rate of 24 frames per second, resulting in a 2-second video. We rendered the animations at a resolution of 1024 × 1024 pixels. In all animations, the ground was set to invisible, and objects were presented against a solid black background. We wanted to ensure that participants do not form vivid impressions or associations of exemplars with any specific object to minimize the likelihood of recalling properties solely based on the object presented in a single frame. Therefore, we deliberately kept the shapes of the objects as generic as possible, such as cubes and spheres, to prevent them from resembling any specific or recognizable objects based on their geometry.

We used Blender version 3.3.0 (Blender Online Community, 2024), an open-source 3D computer graphics software tool, to create animations for our study. We utilized Blender’s different simulation systems to achieve realistic effects across various materials and their interactions with external forces. Details about the rendering of each exemplar from all categories can be found in <u>Appendix A</u>. We rendered three versions of each animation, as described next.

Rendering conditions:

***1. Line:*** To create a line drawing version of animations, we used the grease pencil toolbox within Blender, enabling us to render only occluding contours and creases. A grease pencil object was created, and a line art modifier was added to trace the contours of the selected object. Most settings remained at default, with the line color set to white. Line thickness was adjusted as a function of distance from the camera, maintaining a ratio of line thickness to camera distance between 2 and 3. This ensured uniform thickness despite different camera specifications across exemplars. The final animations rendered only the Grease Pencil outlines (occluding contours and creases), without materials or background, using Blender’s Cycles render engine. Representative frames for one exemplar from each category are shown in the left column of Figure 4.
***2. Dot:*** To generate dynamic dot versions of animations, we imported 3D vertices of each material’s mesh from Blender into MATLAB (version 2022a, MathWorks Inc., New York, NY, USA) and computed 2D projections of vertices based on the camera angles used to render animations. We sampled a subset of 2D vertices and plotted a white dot on a black background for each vertex, frame by frame, to render an animation representing the motion of material through the dots. The number of sampled vertices varied across exemplars based on the area spanned by the material across frames. We first converted the fully textured animations to binary format and calculated the area spanned in each frame as the ratio of pixels with intensity 1 to those with intensity 0. To determine the number of sampled vertices, we used the 75th percentile of the distribution of the area spanned in each frame and multiplied it by 2000. The 75th percentile was chosen instead of the maximum area to avoid outliers, as the distribution of the area spanned across frames is not always normal. Some videos exhibit highly skewed distributions, where using the maximum area would lead to extreme values (See Supplementary Figure S1). By taking the 75th percentile, we ensured a relatively more stable and representative measure. Representative frames for one of the exemplars from each category are shown in the right column of Figure 4, along with corresponding frames from the full condition in the middle column.
***3. Full:*** Full-textured versions of animations (shown in Figure 3) were rendered with all cues, including optical, shape, and motion cues, using Blender’s Cycles render engine. In the remainder of this article, we will refer to it as the *‘full’* condition.

**Figure 4.**
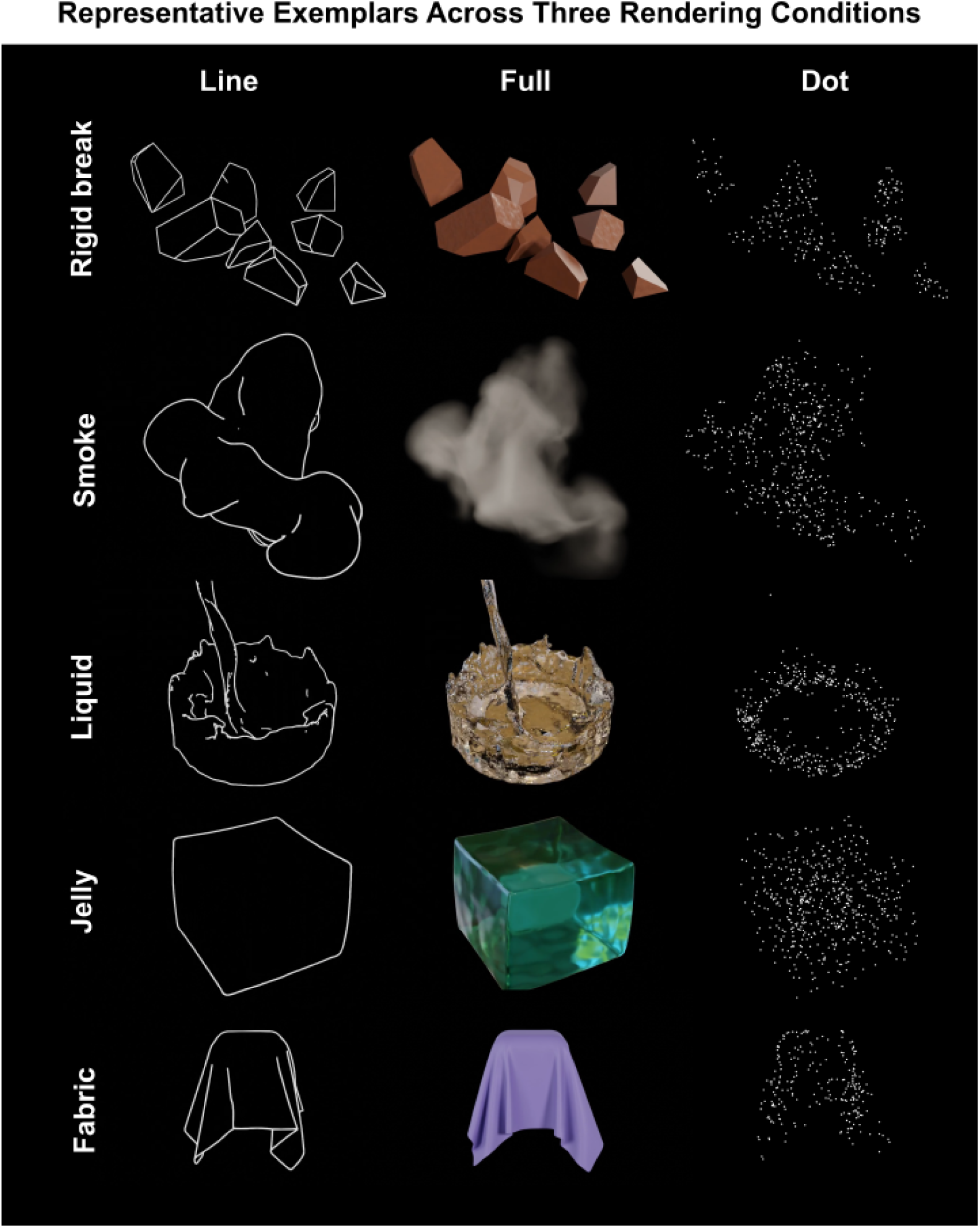
Rendering conditions for stimuli: Each column represents one of the three rendered versions of stimuli, showing a single frame from a representative exemplar for each material category

### Experiment 1: Rating mechanical properties

In the first experiment, participants rated each stimulus on five mechanical material attributes. The experiment followed a between-subjects design with three rendering conditions: line, dot, and full, and 20 different observers participated in each condition. This design was used to prevent participants in the line or dot conditions from recognizing materials by referencing the full condition.

#### Participants

Sixty participants (12 male and 48 female, age range: 19-52 years; mean age: 26.25 years) participated in the experiment. All participants had normal or corrected-to-normal vision (self-report). Participants gave their written informed consent before the experiments. Experimental procedures were approved by the ethics board at Justus-Liebig-University Giessen and were carried out in accordance with the guidelines set forth by the Declaration of Helsinki. Participants were compensated at the rate of 8 euros per hour.

#### Attributes

Initially, we selected eight attributes conceptually divided into two categories: high-level attributes (dense, flexible, wobbly, fluid, and airy motion) to characterize the mechanical qualities of five material categories and low-level attributes (motion coherence, oscillatory motion, and motion dynamics) to relate them to measured motion characteristics of the stimuli, and to assess their contributions to perceived similarity between material categories. However, during a first data collection phase (N = 27), nearly all participants reported difficulty with making judgments on these low-level attributes. A brief inspection of the corresponding rating data revealed that the response distributions for these attributes were nearly uniform across all categories, consistent with participants’ reports of finding it hard to rate stimuli on these attributes. Based on this, we decided to exclude low-level attributes for the remainder of the data collection. The remaining 33 participants were asked to rate only the five high-level attributes on the scale from *0* to *1*. Data on the five high-level ratings were combined across the two groups of participants. Before the experiment, handouts were provided to ensure clarity, outlining each attribute and specifying the criteria for scores *0* and *1* (Figure 5). A translated handout was available for German speakers, but the experiment was always presented in English. The following definitions in the handouts explained the five attributes and their ratings:

**Figure 5.**
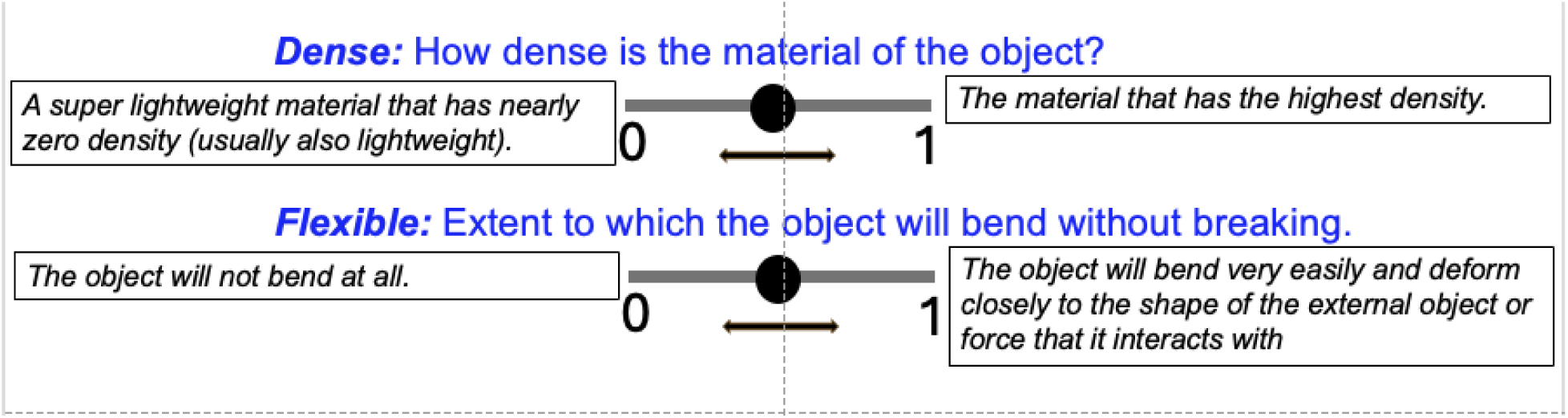
Example of the rating slider used in the experiment, with criteria for scores 0 (left) and 1 (right) provided separately in participant handouts.

***Dense:*** How dense is the material of the object?

**0**: A super lightweight material that has nearly zero density (usually also lightweight).

**1**: The material that has the highest density.

***Flexible:*** The extent to which the object will bend without breaking.

**0**: The object will not bend at all.

**1**: The object will bend very easily and deform closely to the shape of the external object or force that it interacts with.

***Wobbly:*** The extent to which the object will shake or jiggle unsteadily while remaining intact.

**0**: The object will not wobble at all.

**1**: The object will wobble nonstop when triggered by a very subtle force.

***Fluid:*** The extent to which the object will flow.

**0**: The object will not flow at all.

**1**: The object will flow very easily and quickly.

***Airy motion:*** The extent to which the material floats and/or spreads in the air in a lightweight fashion.

**0**: The motion is not airy at all.

**1**: The motion is extremely airy.

#### Experimental setup

Stimuli were presented on a 27-inch LCD monitor (ViewSonic VA270-H, Walnut, California, USA) with a refresh rate of 100 Hz and a resolution of 1920 × 1080, controlled by a Dell system running on Windows 10 Pro. The experiment was developed using a combination of HTML and JavaScript and hosted on a local server. We used the JavaScript library p5.js to present our stimuli. The viewing distance was fixed at 70 cm. The experiment took place in a dark room.

#### Procedure

Upon arrival, participants signed the consent form and received detailed instructions outlining the task and attributes as described above. The experiment started with a welcome screen, and the subject ID was recorded. By pressing the ‘Enter’ button, they proceed to the experimental trials. On each trial, participants viewed the animation of material on the left side of the display, projecting a viewing angle of 18.8^°^. Five sliders corresponding to five material attributes were displayed on the right side of the display with a ‘submit’ button at the bottom (see left panel of Figure 6). Participants were instructed to rate each attribute on a scale from 0 (left) to 1 (right) by dragging the slider with a mouse. The duration of each animation was 2 seconds (24 frames/s), which was replayed continuously until the participants completed rating five attributes and clicked the submit button to proceed to the next trial. The subsequent trial started with an interval of 300 ms. The position of the sliders is reset to the middle of the bar at the start of each trial. There was no time limit, but participants were encouraged not to spend more than 10 seconds on each trial. On average, they spent 20.4 seconds on one trial. The experiment consisted of 114 trials, with one animation per trial. Each unique stimulus was presented only once without repetition, and the order of presentation was randomized. The trial and block numbers were displayed on the bottom right and left corners, respectively.

**Figure 6.**
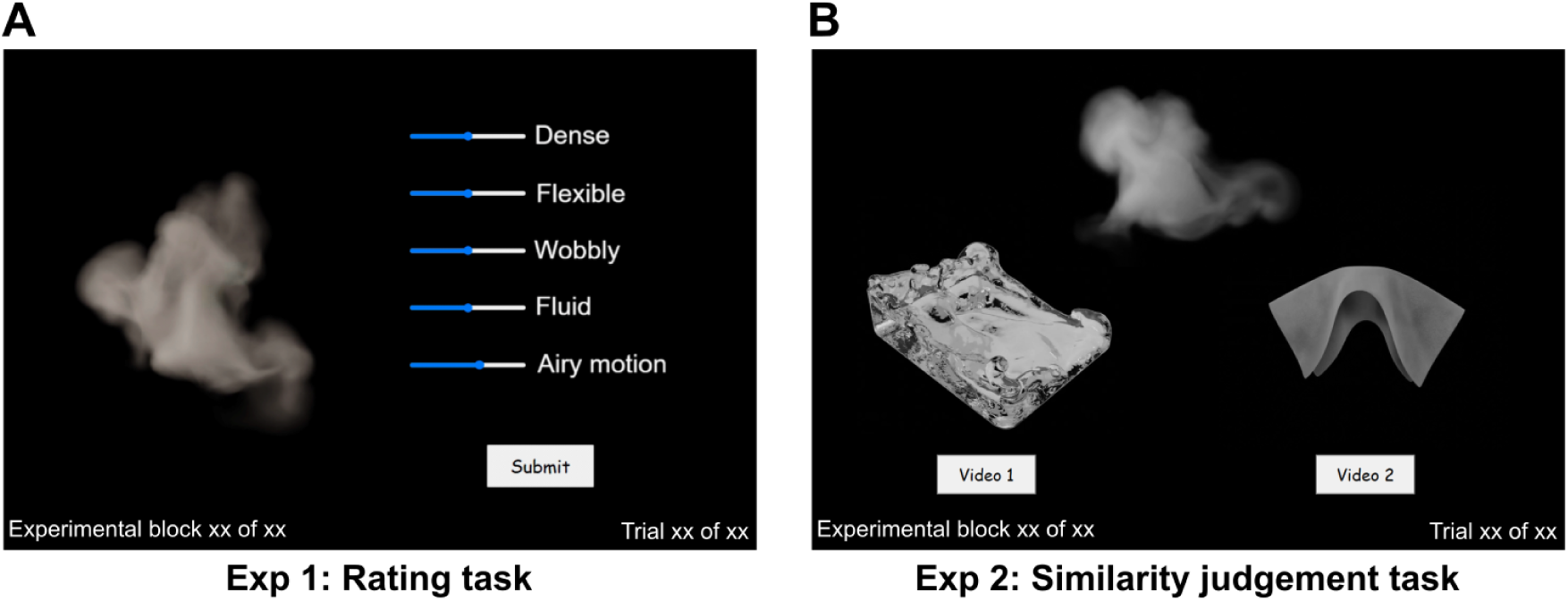
An example of trial layout from A) Experiment 1 - Rating task, and B) Experiment 2 - Similarity judgment task. The text and images have been rescaled in the figure for clarity and do not reflect the actual scale used in the experiment.

#### Data analysis

All analyses and plotting, except cluster analysis, were conducted using RStudio version 2023.9.1.494 (RStudio Team, 2023).

##### Inter-observer correlations

To assess the consistency of attribute ratings across participants, we computed inter-observer correlations. The ratings for all 114 movies (5 attributes per movie) from a single participant were flattened into a vector. The Spearman correlation between these vectors was estimated for each pair of participants, providing a single correlation coefficient per participant pair.

##### Cluster analysis

We analyzed the rating data with a K-Means clustering algorithm. K-Means clustering was implemented using Python’s Scikit-learn library (Pedregosa et al., 2011). We fixed the number of clusters at 5, based on the prior knowledge of five material categories in our data. The centroids were initialized using the K-means++ algorithm, and the algorithm ran until convergence with a default tolerance of 1 x 10^-4^ was reached. The effectiveness of clustering was evaluated using the Silhouette Score and by examining the distribution of trials from different material categories within each cluster. To visualize the results, the trial count was plotted for each cluster, and cluster centroids were visualized in a five-dimensional space representing five material attributes being rated.

##### Representational similarity analysis

To obtain Representational Dissimilarity matrices (RDMs) showing dissimilarities between different materials, we calculated a ‘1 - Spearman’ correlation between the ratings for each pair of exemplars across all participants. Correspondence between RDMs from different rendering conditions was assessed using the Mantel test. For each comparison, Mantel’s *r* indicating Spearman’s correlation between the RDMs was computed, along with a p-value (adjusted for multiple comparisons) based on 999 permutations to assess statistical significance.

### Experiment 2: Similarity judgement task

In the second experiment, we used a similarity judgment task in a triplet two-alternative forced-choice (2-AFC) setting to measure the perceived dissimilarities between materials using the same set of stimuli as in Experiment 1 (Rating task), without requiring participants to rely on semantic labels. The only difference was that we used black-and-white animations for the full condition to avoid color as a strong cue for grouping materials together. The experiment followed a between-subjects design, with three rendering conditions: line, dot, and full, each presented to a separate group of participants.

#### Participants

We recruited 100 participants (no exclusions; 32 male, 67 female, one other; age range: 19–38 years; mean age: 26.25 years). Before the experiment, participants gave written informed consent. Experimental procedures were approved by the ethics board at Justus-Liebig-University Giessen and were carried out in accordance with the guidelines set forth by the Declaration of Helsinki. Participants were compensated at the rate of 8 euros per hour.

#### Experimental setup

This experiment was conducted online, hosted on Amazon S3, and developed using HTML and JavaScript. We used the JavaScript library p5.js to present our stimuli (McCarthy et al., 2015). Participants completed the experiment using a web browser on their personal computers, with the recommended browser being Google Chrome. We instructed participants not to use mobile phones or tablets for the experiment. We offered participants the option to complete the experiment in either English or German, allowing them to choose their preferred language.

#### Procedure

Upon accessing the website, the experiment started with a *welcome screen* that provided general instructions about the online experiment and also directed participants to the informed consent. Next, participants completed a screen calibration procedure (based on the method developed by Li et al. (2020); see Appendix B for details). After calibrating their screens, participants filled out a *demographic form*, providing only age and gender information. They were then provided with information about the experiment and task on an *instruction screen* before the experiment began. They entered the experiment by clicking an ‘Enter’ button at the bottom of the instruction screen. On each trial, participants viewed the three animations, each belonging to a different material category. Animations were displayed in a triangular setting, as shown in the right panel of Figure 6. The size of the animations was adjusted according to viewing distance such that each animation projected a viewing angle of 10^°^. The animation presented at the top was designated as the *reference* stimulus, while the two shown at the bottom served as *test* stimuli. Participants were asked to choose which of the two test animations was more similar to the reference animation, based on the perceived material. They indicated their choice by clicking the button located below the selected animation. The buttons were outlined in red from the beginning of the trial until the videos played once, i.e., for 2000 ms. We instructed participants not to respond until the outline disappeared. Even if they tried to respond before that, the subsequent trial would not start, forcing them to view the animations at least once. There was no time limit, but they were encouraged not to spend more than 10 seconds on each trial. On average, they spent 5.6 seconds on one trial. After they responded, the subsequent trial started with an interval of 300 ms. We also included catch trials to ensure participants maintained their focus throughout the experiment. If a participant consistently chose the same-sided video for more than 14 consecutive trials, or if their reaction times (measured from the time after the red line from the response buttons disappeared) were consistently shorter than 350ms, they were presented with a catch trial. During catch trials, participants were shown a warning message: ‘Are you paying attention?’ followed by an unrelated task (e.g., counting the number of circles among different shapes) to re-engage their attention.

#### Construction of representational dissimilarity matrices (RDMs)

To derive the perceived similarity between material categories, we followed the approach of Schmidt et al. (2025), where RDMs are constructed from participants’ similarity judgments in a triplet-based task. These judgments are aggregated across all triplet contexts to compute pairwise similarity scores, which are then transformed into dissimilarity values (see Appendix C for full details).

Correspondence between rendering conditions was assessed by comparing RDMs across different rendering conditions within the similarity judgment task. Additionally, to assess cross-task correspondence, RDMs from the similarity judgment task were compared with those from the rating task. Both comparisons were evaluated using the Mantel test. For each comparison, Mantel’s *r* indicating the Spearman’s correlation between the RDMs was computed, along with a p-value (adjusted for multiple comparisons) based on 999 permutations to assess statistical significance. All analyses and plotting were conducted using RStudio version 2023.9.1.494 (RStudio Team, 2023).

### Motion and shape statistics

To identify features critical for estimating material properties, we analyzed motion and shape statistics derived from full-textured animations. We computed 20 motion statistics following the methodology of Kawabe et al. (2015) and 20 shape statistics following the methodology of Paulun et al. (2015). Motion statistics comprised four statistical moments, mean, standard deviation (SD), skewness, and kurtosis, computed for five distinct image-based motion features: image speed, divergence, curl, gradient, and discrete Laplacian. These calculations were implemented in MATLAB using vertices projected onto 2D frames (refer to Kawabe et al. (2015) for details). We computed a set of 20 shape-related features capturing aspects such as curvature, orientation, spatial distribution, and overall geometry of the objects in the animation frames (refer to Paulun et al. (2015) for details). Shape computations utilized binary masks obtained from the projected 2D animation frames and were likewise implemented in MATLAB. For each of the 114 animations, every one of the 40 total features (20 motion + 20 shape) was represented by a 48-dimensional vector, with each dimension corresponding to a single animation frame.

The analysis was done separately on shape and motion statistics. First, each feature column was transformed into z-scores so that all features contribute equally to downstream analyses. We then performed principal component analysis (PCA), retaining the minimal number of components that explained at least 80% of the variance. We then aggregated the resulting PC scores by computing the mean score of each retained component across all 48 frames of each video, yielding one representative PC score vector per video. These representative vectors defined each video’s coordinates in the PCA space. Next, to predict each of the five attributes (dense, flexible, wobbly, fluid, and airy), we identified a reference video that had received the lowest rating for the target attribute in the rating experiment (Experiment 1). We then computed the Euclidean distance from this reference video to all other videos in the PCA space, thereby yielding a predicted attribute value for each video based on its distance to the reference.

To evaluate how well motion and shape statistics predicted each of the five perceptual attributes, we conducted linear regression analyses relating the predicted attributes to the perceptual attributes (from Experiment 1). Adjusted R-squared values were extracted to assess model fit, and p-values were computed to determine statistical significance. To maintain reliable statistical conclusions, we employed the Benjamini-Hochberg procedure for multiple comparison corrections, thereby controlling the false discovery rate.

In order to assess how well motion and shape statistics account for similarity judgements in Experiment 2, we constructed dissimilarity matrices based on the predicted attributes using the same methodology applied to the perceptual data. These RDMs were compared to the perceptual RDMs derived from the rating and similarity judgment tasks using the Mantel test. For each comparison, Mantel’s *r*, reflecting the Spearman correlation between RDMs, was computed alongside permutation-based p-values (adjusted for multiple comparisons), using 999 permutations to assess significance. All analyses and visualizations were performed using RStudio version 2023.9.1.494 (RStudio Team, 2023).

## Results

### Experiment 1: Rating task

The purpose of Experiment 1 was to assess whether participants are able to perceive mechanical qualities of materials based on contour motion alone (line drawing condition), and to compare their performance to that obtained with dynamic dot and full-textured animations. Participants viewed animations of materials and rated five mechanical attributes: density, flexible, wobbly, fluid, and airy motion, which were chosen to capture the characteristic features of the material categories used in our study. One of the common issues associated with the use of rating scales is the central tendency bias, where participants tend to avoid the extreme ends of the scale and prefer responses closer to the midpoint of the scale, leading to regression to the mean (Douven, 2018; Stevens, 1971). This bias can distort data by underrepresenting extreme values of responses. Given this concern, we first examined the distribution of our rating data. Figure 7 shows the rating histogram pooling values across all participants, attributes, material categories, and conditions. This shows a bimodal distribution with clustering of responses to extreme values (0 and 1) rather than intermediate values. The frequent use of extreme values suggests that participants have strong categorical impressions of the perceptual qualities being evaluated. Thus, central tendency bias does not appear to be present in our data.

**Figure 7.**
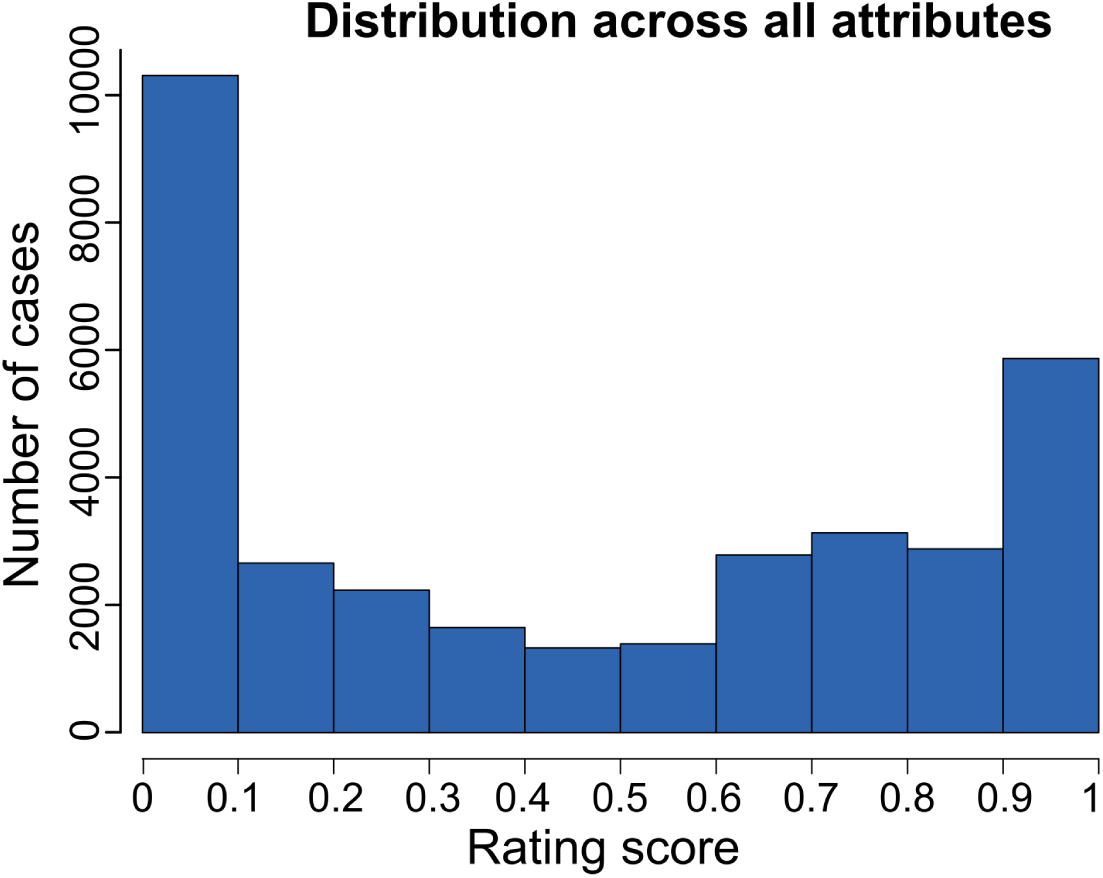
Distribution of attribute rating values pooled across all participants, attributes, material categories, and conditions.

We also noted that the peak at the lower extreme is notably larger. While this asymmetry could suggest a potential response bias toward low values, it is more plausibly explained by the stimuli’s inherent characteristics. All the attributes in the rigid-breakable category except ‘dense’ naturally cluster around lower extreme values, contributing to this imbalance, as can be seen in the mean ratings of the rigid-breakable category in Figure 8. Each material displays a unique characteristic pattern, with ratings spanning a range of values from high through intermediate to low, thus reflecting the distinct perceptual properties associated with each category.

**Figure 8.**
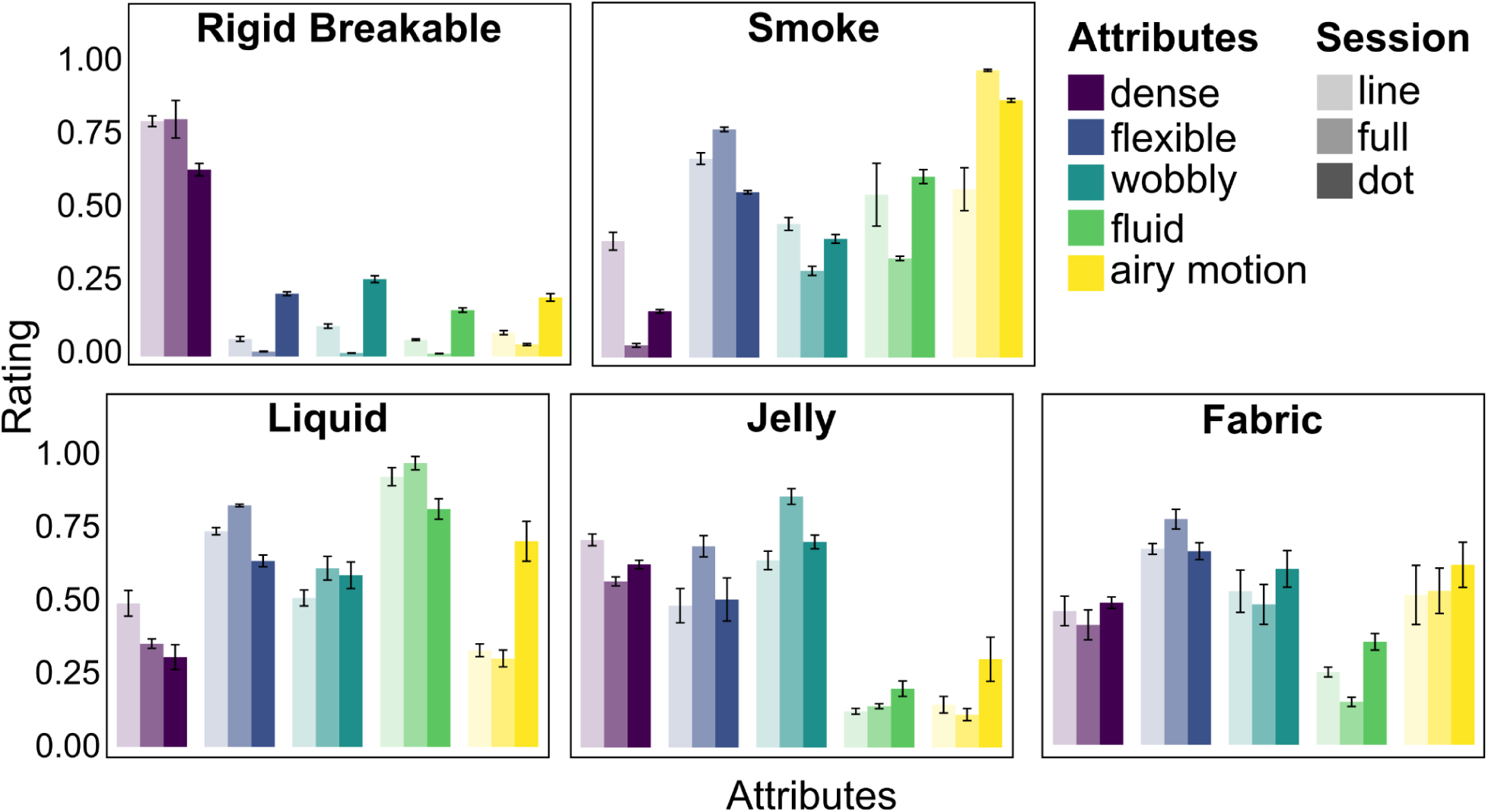
The averaged ratings for each attribute across materials and three rendering conditions are represented by bar intensities corresponding to alpha values. The bars with the lowest saturation reflect ratings from the ‘line’ condition, those with medium saturation correspond to the ‘full’ condition, and the highest saturation represents ratings from the ‘dot’ condition. We present the results from the ‘full’ condition between ‘line’ and ‘dot’, in order to allow for better visual comparison.

#### Inter-observer correlations

We examined inter-observer correlations to determine whether participants were consistent in their ratings. If there is inconsistency, it suggests that the selected attributes are interpreted subjectively, potentially lacking clarity or shared meaning. Conversely, if participants are consistent in their ratings, it suggests that the attributes are meaningful and objective.

Figure 9 shows the Spearman correlation between participants for all three rendering conditions. The mean inter-observer correlations for the line, full, and dot conditions are 0.75, 0.82, and 0.66, respectively. All inter-observer correlations were substantially positive for all conditions, ranging from 0.43 to 0.93, and were significant at the level of *p* < 0.001. The mean inter-observer correlation for the line and dot conditions was lower than for the full condition. However, considering that these were significantly reduced conditions of fully textured animations, the correlations remain high. Overall, the high inter-observer correlations for all conditions indicate that the selected attributes were meaningful and objective, effectively capturing the characteristic material properties across different categories. Another observation was that the inter-observer correlations were higher for the line condition compared to the dot condition, which indicates that the absence of clear contour cues leads to more ambiguity in inferring the mechanical attributes.

**Figure 9.**
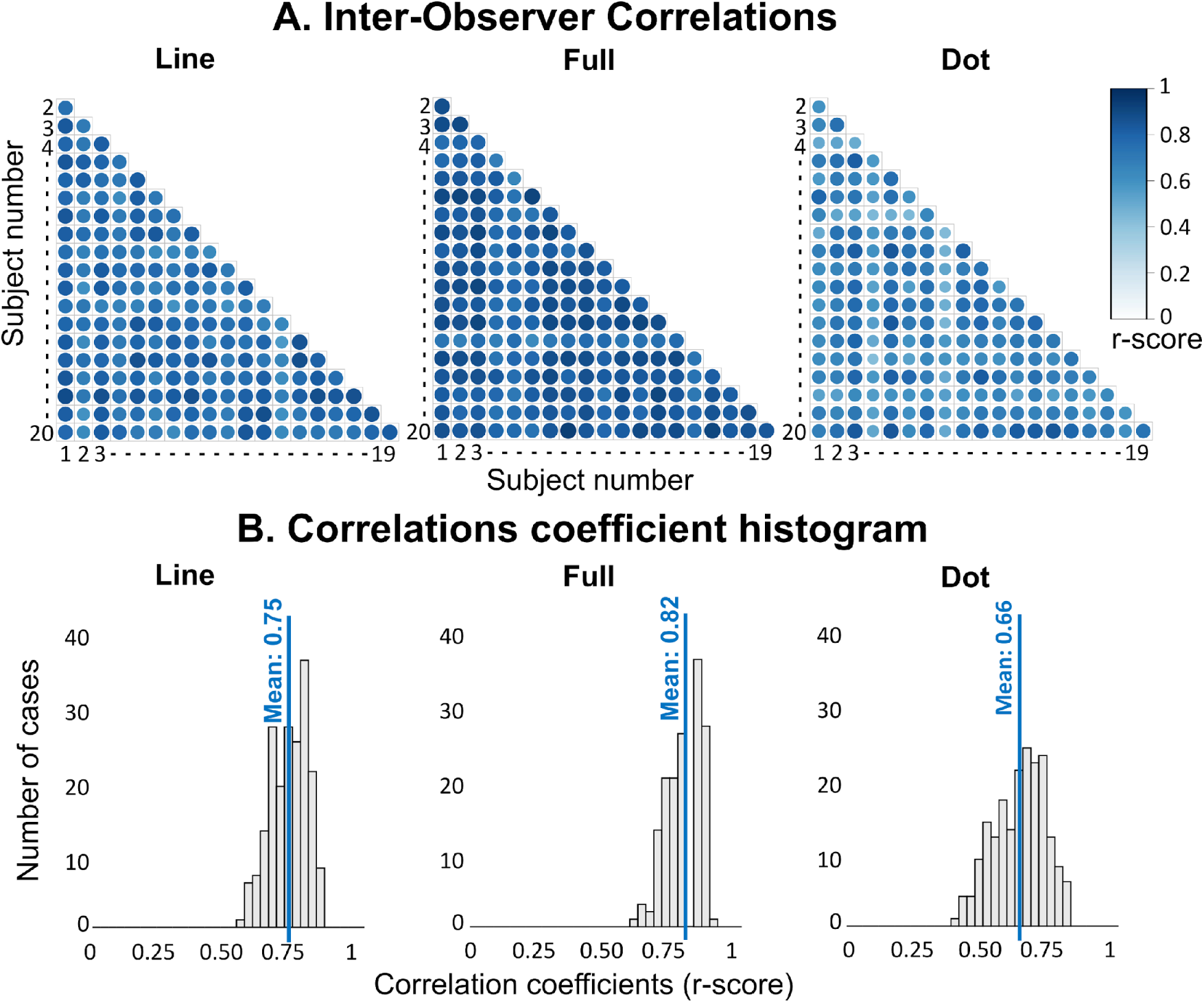
Spearman correlations between participants in Experiment 1. (A) Inter-observer correlations (i.e., correlations between each participant and all other participants) for line, full, and dot conditions. Correlation coefficient values (r-scores) are represented by the color gradient shown by the bar on the right side. A lighter shade of blue indicates a lower correlation, while a darker shade indicates a higher correlation. All the correlations are significant at the level of p<0.001 and positive, ranging from 0.43 to 0.93. (B) Histogram of the correlation coefficients between all 20 participants across all stimuli and attributes tested for line, full, and dot conditions, respectively, from left to right. Mean correlations are labeled by a blue vertical line for each condition.

#### Cluster analysis

We performed cluster analysis on the rating data for each rendering condition, line, full, and dot, to compare how material qualities are perceived across these conditions. Figure 10-A shows the results of k-means clustering with 5 clusters. The clustering quality was assessed using the Silhouette Score, with the *‘full’* condition serving as the baseline for comparison. Each cluster was labeled based on the material category that contributed the highest trial count. The centroids of these representative clusters were plotted in a five-dimensional space, reflecting the five perceptual attributes associated with each labeled material category, as shown in Figure 10-B. Each row corresponds to the cluster centroids for one of the three rendering conditions: line, full, and dot, with the labeled categories assigned to the clusters shown on the top row.

**Figure 10.**
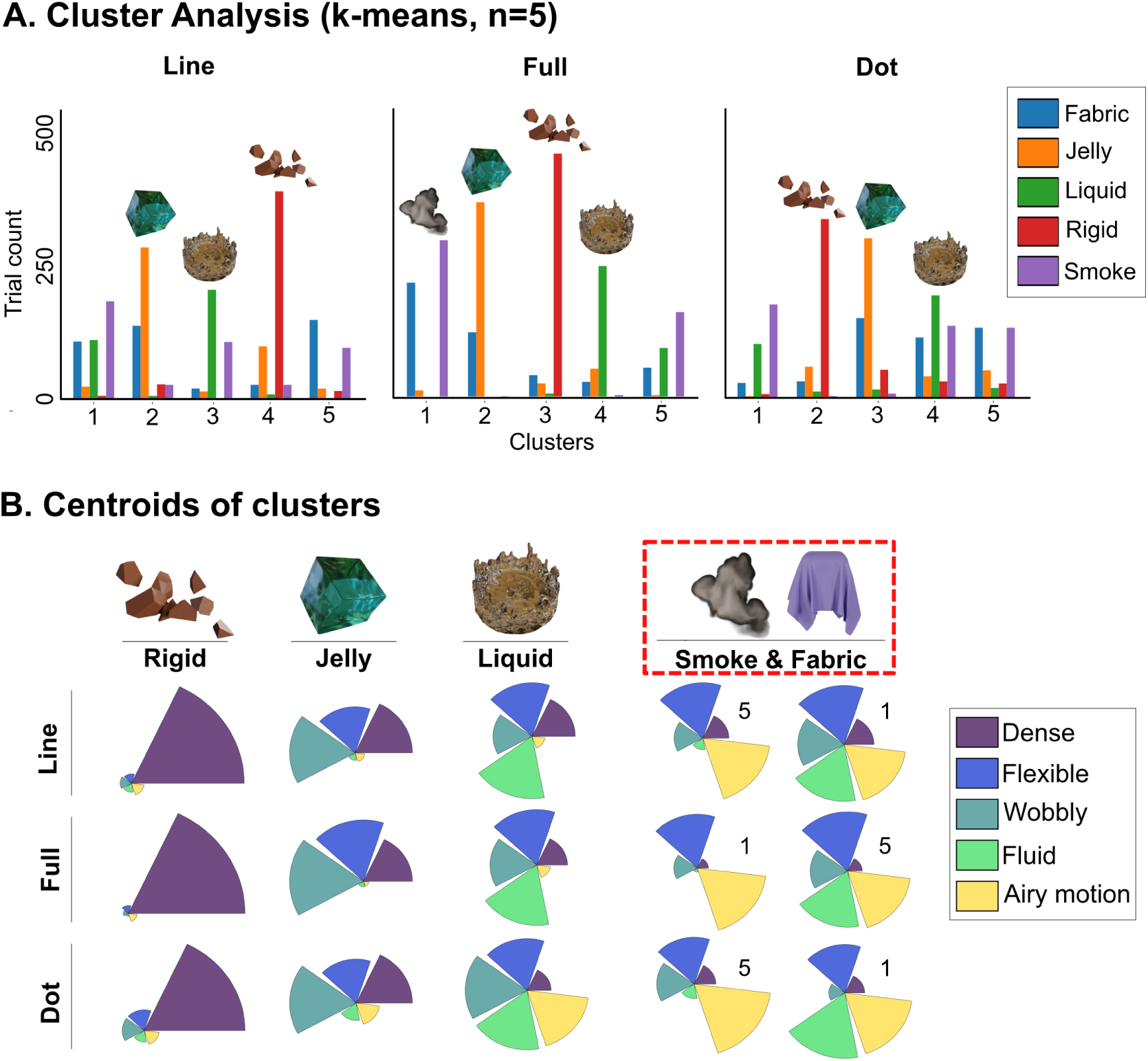
Cluster analysis with k-means clustering (n=5) (A) Cluster distributions from the clustering analysis for the line, full, and dot conditions. The y-axis represents the trial count, and the x-axis displays the five clusters. Each cluster contains multiple bars, color-coded to indicate trials from different material categories. The category with the highest trial count in each cluster is designated as the representative category, shown with a material thumbnail above the cluster. (B) Centroids of the clusters from panel A are plotted in a five-dimensional space, with each attribute represented by a specific color. The representative category is displayed at the top of each cluster. The three rows, from top to bottom, correspond to the three rendering conditions: line, full, and dot. The last two centroids, highlighted with a dotted rectangle on top, indicate mixed clusters without a single dominant category. The number on the centroid corresponds to the cluster number in Panel A, representing the distribution of categories in these clusters.

***Full:*** The cluster distribution from the full condition is shown in the middle column of Figure 10-A, which achieved a Silhouette Score of 0.38, indicating moderately good clustering performance. Three clusters were distinctly dominated by trials from the rigid-breakable, jelly, and liquid categories, showing a clear separation for these groups. However, the overall moderate score reflects partial overlap between the fabric and smoke categories, indicating less distinct boundaries and reduced cluster separation in these cases. The middle row of Figure 10-B displays the cluster centroids for each category from the full condition, representing the characteristic properties of each material type.

***Line:*** The line condition achieved a Silhouette Score of 0.33, slightly lower than the 0.38 observed for the full condition, but the clustering structure remained largely intact. The centroid distributions and attribute proportions for categories like fabric, jelly, and rigid-breakable showed strong alignment with those in the full condition, indicating minimal perceptual divergence for these categories. However, a notable difference was observed in the line condition, where liquid and smoke shared more perceptual space than in the full condition.

***Dot:*** The *dot* condition scored the lowest Silhouette Score of 0.27, reflecting weaker clustering quality compared to the full condition. This weaker performance is attributed to the increase in shared perceptual space among liquid, smoke, and fabric categories, where clusters other than clusters 2 and 3 lack a distinctly dominant category. A comparison of the cluster centroids between the *dot* and *full* conditions reveals that while the characteristic attributes of the rigid-breakable and jelly clusters remained largely consistent, some subtle variations emerged. Notably, in the liquid cluster, attributes such as airy motion and wobbliness appeared slightly more pronounced.

Overall, while both the dot and line conditions exhibited similarities to the full condition, the line condition was more similar to the full condition in terms of clustering patterns, suggesting that contour motion preserves some key perceptual characteristics of materials better than dynamic dot materials.

#### Representational similarity analysis

RDMs across three rendering conditions: line, full, and dot, are shown in the upper row of Figure 11, depicting the perceived differences between five material categories. Upon visual inspection of the RDM from the full condition (upper middle panel of Figure 11), a clear pattern of perceptual differences emerges. Materials belonging to the same category were perceived as highly similar to each other while being more dissimilar to materials from other categories. Furthermore, the perceptual differences between categories appear consistent across exemplars within those categories. For instance, the rigid-breakable category is perceived as highly dissimilar to the liquid category, and this dissimilarity pattern holds consistently for all exemplars of liquid and rigid-breakable materials. This shows that the mechanical attributes we chose are characteristic properties of the material category and did not vary within the category, despite varied motion patterns within a category.

**Figure 11.**
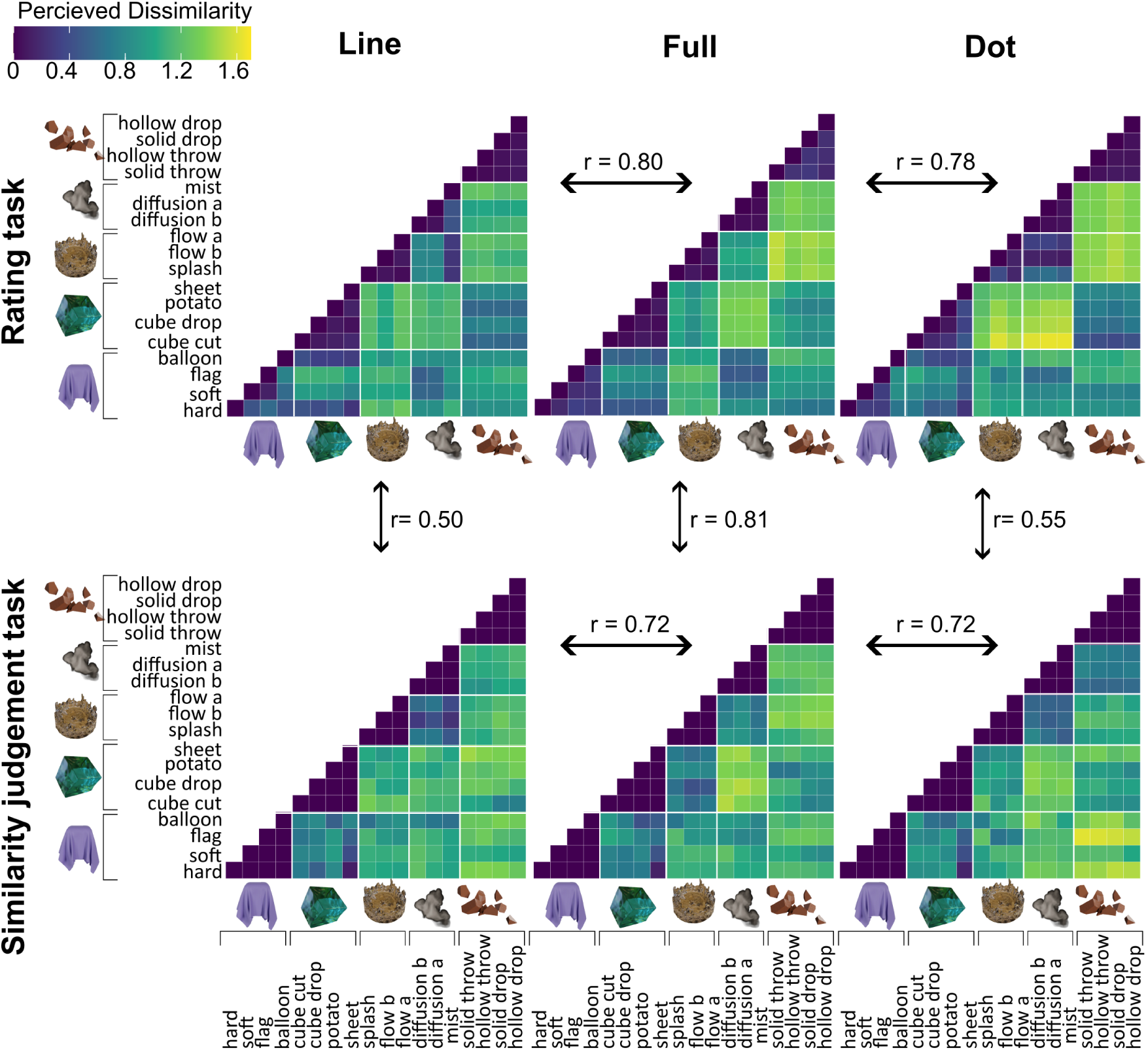
Representational Dissimilarity Matrices (RDMs) illustrating perceived similarities between material categories across three rendering conditions—line, full, and dot. The upper row displays RDMs derived from the rating task, while the lower row shows RDMs from the similarity judgment task. Arrows indicate the Mantel test correlations (r) between corresponding RDMs across tasks and rendering conditions

Next, to see whether these perceptual similarities are preserved when optical cues are removed, we compared the RDMs obtained from the line and dot conditions with the full condition. After correcting for multiple comparisons (*p* < 0.007), the Mantel test showed a strong and statistically significant relationship. Specifically, the Mantel test statistic (*r*) for the comparison between the full and line RDMs was *r* = 0.802, *p* = 0.001, and for the comparison between the full and dot RDMs, it was *r* = 0.784, *p* = 0.001. This correspondence is also visually evident in the patterns observed within the RDMs. However, some differences between line and dot conditions were apparent, which may offer insights into the relative importance of internal motion and contour motion cues in the perception of these material categories. Overall, the results strongly suggest that both line and dot conditions effectively convey the mechanical properties of materials.

### Experiment 2: Similarity judgment task

To validate whether the five-dimensional space in Experiment 1 sufficiently characterizes the set of materials, and to ensure that perceptual dissimilarity measures were not biased by the use of a verbal task, we conducted an additional experiment. In this experiment, we constructed representational dissimilarity matrices (RDMs) based on non-verbal similarity judgments obtained through a triplet 2-alternative forced-choice (2-AFC) task. The lower row in Figure 11 shows the RDMs derived from this experiment.

Visual inspection of the RDMs from the full condition of experiments 1 and 2 revealed consistent patterns between the rating and similarity tasks (see the top middle columns of Figure 11 compared to the lower middle column). This correspondence is supported by very high r statistics derived from the Mental tests (*r* = 0.805, *p* = 0.001). Additionally, the correlation between the RDMs of line and dot conditions across task types is also significant (dot rating RDM and dot similarity judgment RDM: *r* = 0.555, *p* = 0.001; line rating RDM and line similarity judgment RDM: *r* = 0.497, *p* = 0.001), albeit lower than that observed for the full condition. These results suggest that the rating attributes we chose captured the perceptual space of materials fairly well, and that it is not biased by having to make verbal responses.

Further, results indicate a statistically significant correlation between RDMs from line, dot, and full conditions (lower panel of Figure 11). The Mantel test statistic (*r*) comparing the line and full RDMs is *r* = 0.721, *p* = 0.001, while the comparison between dot and full condition yielded r = 0.718, *p* = 0.001. These results align with the results from experiment 1, indicating that participants perceive materials in the line and dot conditions similarly to the full condition. The RDMs derived from the similarity judgment task revealed slightly greater differences between reduced (line and dot) and full conditions compared to those from the rating task. This suggests that when grouping materials by visual similarity, participants consider additional dimensions not captured in our rating experiment. We discuss this further below.

### Motion and shape statistics

Figure 12-A shows the results of the regression analysis relating 20 motion and 20 shape features (described in the analysis), each to the 5 rated material attributes. Results from motion statistics suggest that *fluid* had the highest explanatory power (adjusted *R^2^* = 0.36), followed by *airy motion* (adjusted *R^2^*= 0.26), *flexible* (adjusted *R^2^* = 0.25), *dense* (adjusted *R^2^* = 0.09), and *wobbly* (adjusted *R^2^*= 0.03). In contrast, shape-derived statistics demonstrated stronger predictive power for *wobbly* (adjusted *R^2^* = 0.47) and *flexible* (adjusted *R^2^*= 0.42), with lower values for *dense* (adjusted *R^2^* = 0.11), *fluid* (adjusted *R^2^* = 0.07), and *airy motion* (adjusted *R^2^* = 0.02). These findings suggest that motion statistics and shape statistics are each predictive of different material attributes, and neither of them appears to be informative for predicting perceived density. We next conducted Mantel tests to compare RDMs derived from motion features with perceptual RDMs obtained through rating tasks and similarity judgments (Figure 12). For the rating RDMs, we observed small-to-moderate correlations (Mantel’s *r* = 0.256 – 0.403) that remained significant after multiple-comparison corrections (*p_corr_* = 0.012 – 0.036). Notably, the line condition showed the highest correlation among the rating RDMs. By contrast, correlations with the similarity judgement RDMs were small (Mantel’s *r* = 0.104 – 0.244) and were not significant. These findings indicate that motion-derived features capture participants’ attribute rating judgments more effectively than their similarity judgments.

**Figure 12.**
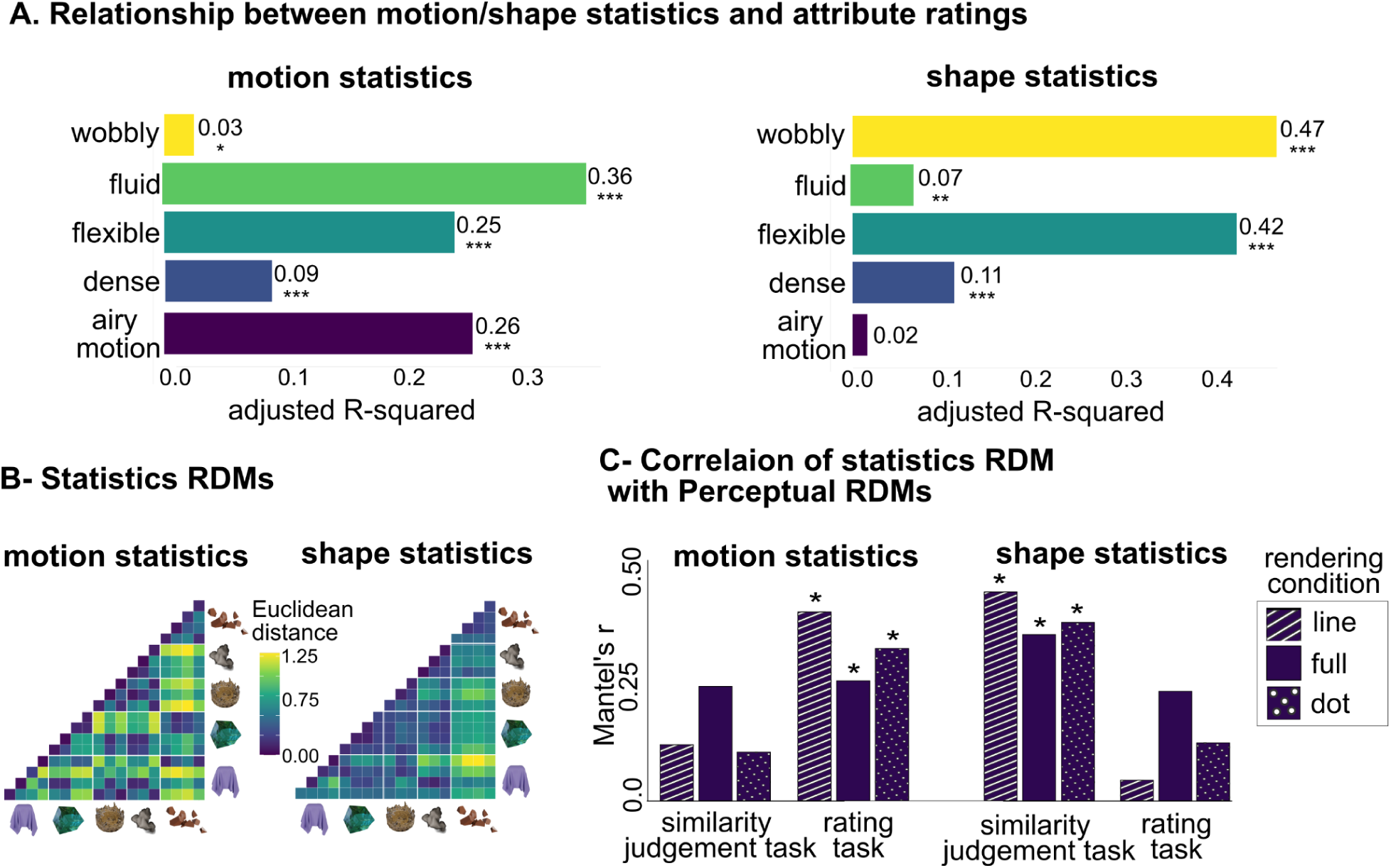
Relationship between perceptual attributes and underlying computational representations based on motion and shape features. (A) Adjusted R2 values from regression analyses predicting perceptual attributes with motion and shape statistics (* for p<0.05, ** for p<0.01, and *** for p<0.001, corrected for multiple comparisons). (B) RDMs derived from motion and shape statistics. (C) Mantel’s r values comparing statistics RDMs to perceptual RDMs (rating and similarity judgments) across the line, full and dot conditions (* for p<0.05, corrected for multiple comparisons).

In contrast, shape derived RDMs exhibited the opposite pattern. For the rating RDMs, correlations were small (Mantel’s *r* = 0.043 – 0.232) and were not significant. However, correlations with the similarity judgment RDMs were moderate (Mantel’s *r* = 0.353 – 0.443) and reached significance in all conditions (*p_corr_* = 0.006), with the line condition again yielding the highest correlation. These findings indicate that shape statistics capture participants’ similarity-based judgments more effectively than their attribute rating judgments.

## Discussion

Although static contours have been shown to contribute to material perception (Pinna & Deiana, 2015), they can be ambiguous in the absence of motion cues as illustrated in Figure 1. To determine the contribution of contour motion cues in material perception, we employed animated line drawings depicting only contour motion and demonstrated that contour motion alone can provide sufficient information for material perception. We tested this by comparing perceptual judgments across three rendering conditions: line (depicting contour motion), full (with fully textured material), and dot (depicting dot motion), in two experiments: a rating experiment and a similarity judgment experiment. Overall, the results showed that participants were able to perceive materials from both line and dot conditions comparably to the full condition, with greater similarity between the line and full condition than between the dot and full condition.

It has long been known that the information preserved in line drawing images serves as a basis for the visual recognition of objects and scenes (Biederman & Ju, 1988; Elder, 2018; Elder & Goldberg, 2002; Greene & Oliva, 2009; Sann & Streri, 2007; Walther et al., 2011). However, it is not yet fully understood why line drawings are so robust. A possible explanation for the effectiveness of line drawings, as suggested by Sayim & Cavanagh (2011), is that they activate neural mechanisms originally adapted for processing natural environments. The primary visual cortex contains neurons specifically attuned to contour orientation, which recognize edges in natural settings rather than uniform regions. Due to their sensitivity to contrast and orientation patterns, these neurons not only detect real-world edges but also respond to drawn lines. Since lines in drawings are generally drawn at the locations where edges would appear in reality, these drawings activate the same perceptual processes as real-world images. Certain edges, such as occluding contours and creases, serve as key shape-defining features that allow the brain to infer the object’s underlying 3D structure from a 2D representation (Koenderink, 1984; Markosian et al., 1997). Hence, lines in line drawings serve as contours that convey the structure of 2D shapes, allowing the brain to infer the underlying 3D form. Moreover, neurons in V1 also detect the local motion of edges, while motion-sensitive areas such as MT integrate these signals to infer global object motion and transformations (Born & Bradley, 2005). Thus, the brain interprets moving lines in dynamic line drawings not as isolated features, but as edges of objects undergoing motion or shape changes.

Thus, our findings suggest that contour motion in dynamic line drawings provides a powerful cue by conveying both shape and motion information simultaneously. This dynamic information helps observers perceive not only shape transformations, such as how a fabric folds or stretches, but also motion characteristics, such as the rhythmic flapping of a flag, the speed of a falling rigid object, or the speed of flow of a liquid. Together, these cues allow the visual system to infer material properties efficiently, aligning with previous work that has demonstrated the importance of both shape and motion cues in material perception (Bi et al., 2019; Bouman et al., 2013; Kawabe et al., 2015; Kawabe & Nishida, 2016; Paulun et al., 2017; Schmid & Doerschner, 2018; Van Assen et al., 2018; van Assen & Fleming, 2016). While prior research has provided important insights into how specific cues predict perceived material properties within a given category, for example, the relationship between flow speed and perceived viscosity of liquids (Kawabe et al., 2015), between several motion statistics and perceived stiffness of fabrics (Bi et al., 2019), or between deformation magnitude and perceived stiffness of jelly (Paulun et al., 2017), these studies have typically focused on single material classes. Materials from different categories can sometimes share similar dynamic properties. For instance, a fabric and a liquid might exhibit similar motion speeds when analyzed using optical flow, yet they differ so much in perceived viscosity. Likewise, a loosely woven cloth and a soft piece of jelly may undergo similar magnitudes of deformation, yet they are perceived to differ markedly in stiffness. So, how does the visual system identify whether an object is a piece of cloth, a stream of liquid, or a piece of jelly in the first place? Can observers distinguish between materials from different categories based on dynamic information alone? And if so, what specific image features does the visual system rely on to make these judgments? Our results suggest that motion conveyed solely through dynamic line drawings provides sufficient information for perception properties across a range of material categories, including jelly, liquid, smoke, fabric, and rigid-breakable materials.

### Perceptual consistency across ratings and similarity judgments

The rating task produced RDMs that closely aligned with those from the similarity judgment task, yielding a remarkably high correlation of 0.81 for the full condition. This result highlights two important implications. First, it has been argued that rating tasks rely on an observer’s ability to use language to describe material properties (Xiao et al., 2016). To address this potential linguistic bias, Experiment 2 employed similarity judgment, which bypassed explicit verbal encoding and provided a more direct perceptual assessment. Notably, while the rating experiment required participants to infer five material properties based on motion, shape, or optical cues, the similarity judgment task offered a purely non-verbal measure of how materials were grouped based on perceived properties. The strong agreement between these two tasks suggests that both verbal (semantic) and non-verbal (visual) judgments are interpreted through a shared underlying perceptual framework. This conclusion aligns with previous research showing a high degree of correspondence between the semantic representations of material categories and the attribute ratings of individual materials (Fleming et al., 2013) and between visual and haptic judgments of material properties (Wiebel et al., 2013). While these studies have primarily provided indirect evidence of a link between verbal (semantic) and perceptual representations, our study offers a direct comparison between the perceptual space derived from verbal attribute ratings and non-verbal visual similarity judgments. Second, despite being limited to only five mechanical attributes, the rating task produced RDMs that are strongly correlated with similarity judgment RDMs. This is in line with findings of Schmid & Doerschner (2018), who demonstrated through factor analyses of material ratings that adjectives cluster around a few core dimensions rather than forming an exhaustive or independent set. They identified key perceptual dimensions such as hydration, fluidness of motion, airiness/density, hardness/softness, and smoothness. The attributes we selected (density, fluidity, flexibility, wobbliness, and airy motion) correspond closely to these dimensions. These findings are further supported by recent large-scale, data-driven work on characterization of material representations (Schmidt et al., 2025), showing that only a set of 36 dimensions can account for the majority of explained variance in human similarity judgments across materials sampled from 200 categories.

Together, these findings suggest that material categorization and similarity are governed by a common, low-dimensional perceptual structure that integrates both semantic and sensory information. Even when observers rate only a subset of mechanical material attributes, the resulting perceptual space is still highly predictive of general visual similarity judgments.

### Contribution of dynamic and optical cues to material perception

Strong correlations between the dot and line RDMs with the full condition RDMs in both rating and similarity judgment tasks (Figure 11) demonstrate that dynamic cues alone carry sufficient information for material perception. However, despite these high correlations, closer inspection reveals small yet meaningful deviations in how materials are perceived when optical cues are absent (line and dot conditions). These subtle deviations emphasize the complementary role optical cues play in the perception of material properties.

For instance, cluster analysis (Figure 10A) shows that in the line condition, there is greater perceptual overlap between smoke and liquid clusters compared to the full condition, which could be due to smoke being perceived as more fluid in the absence of its characteristic optical features. Representational similarity analysis (RSA) also shows increased similarity between liquid and smoke in the line condition compared to the full condition. These perceptual shifts become even more pronounced in the dot condition, where liquids overlap significantly with multiple categories, particularly smoke and fabric. Additionally, rigid materials in the dot condition are perceived as more flexible, wobbly, and airy. These observations suggest two key points: 1) Optical cues complement dynamic motion information and are essential for accurately distinguishing certain material categories. 2) Motion conveyed through lines preserves essential information required for material perception more effectively than motion conveyed through dots.

Importantly, deviations from full condition are more pronounced in similarity judgment tasks than rating tasks, as indicated by slightly lower correlations between full and line conditions (0.72 similarity judgment vs. 0.80 rating) and between full and dot conditions (0.72 similarity judgment vs. 0.78 rating). We also observe notable qualitative changes in the patterns of similarity judgment RDMs when comparing different rendering conditions. For example, rigid breakable materials are perceived as more similar to smoke rather than fabric or jelly in the line and dot conditions, whereas under full conditions, these materials are perceived as more similar to fabric and jelly. Why are these shifts specific to the similarity judgment task and absent in the rating task? We suggest that similarity judgments rely on the interplay of both optical and motion cues to group materials meaningfully. Hence, when optical vues are missing (in line and dot condition), it leads to noticeable shifts in perceived similarity. In contrast, explicit attribute ratings of mechanical attributes (e.g., density or fluidity) rely predominantly on distinct dynamic motion cues that sufficiently convey specific properties independently. These findings are in line with prior research in liquid perception, where shape and motion cues dominate when participants rate liquid viscosity, but optical cues become more influential in a category naming task (van Assen & Fleming, 2016). The category naming task, like our similarity judgment task, directly taps into underlying perceptual space. These results suggest that while dynamic cues play a critical role, they are not sufficient on their own; optical cues are also crucial, especially for capturing dimensions beyond those considered in our rating experiment.

This interpretation is further supported by notably weaker correlations between rating and similarity judgment RDMs in the line (0.50) and dot (0.55) conditions. These lower correlations arise because the removal of optical cues affects the two tasks differently: similarity judgments engage a broader set of perceptual dimensions that are more sensitive to the interplay of available cues, whereas attribute ratings focus on a distinct, limited set of dimensions that can often still be judged accurately using motion cues alone. As a result, when optical information is removed or unavailable, the perceptual space underlying similarity judgments is more substantially altered than that of attribute ratings. This task-specific impact leads to different representational geometries for the same stimuli, reducing the correlation between the resulting RDMs and highlighting how the task demands modulate the influence of optical versus dynamic information on material perception.

### Motion and shape statistics

Previous research has identified several motion and shape statistics that can predict perceived properties of materials such as viscosity in liquids or elasticity in jelly (Kawabe & Nishida, 2016; Paulun et al., 2015, 2015; Van Assen et al., 2018; van Assen & Fleming, 2016). However, a question is: how generalizable are these predictive relationships across different material categories and properties? Our findings suggest that motion statistics moderately predict fluidity, while shape statistics moderately predict attributes such as wobbliness and flexibility across all material categories. These moderate effects suggest that additional factors might be involved in shaping perceptual judgments across material categories.

A comparison of statistical RDMs, derived from motion and shape calculations, with perceptual RDMs revealed distinct patterns. Motion statistics correlated more strongly with the rating task, whereas shape statistics exhibited stronger correlations with the similarity judgment task. Importantly, the highest correlations for motion statistics with rating tasks and shape statistics with similarity judgments were observed in line and dot conditions, with correlations being lowest in the full condition. This suggests that 1) shape information may play a more significant role when participants group materials based on overall similarity. In contrast, when explicitly rating specific material attributes, participants rely more on motion statistics. We suggest that this pattern arises due to the nature of the task. In a similarity-judgment task, where three animations are presented side by side, participants naturally group materials by their overall shape and deformation pattern. Indeed, in a large-scale study using static images of 600 material exemplars, Schmidt et al. (2025) showed that only a limited set of 36 perceptual dimensions underlies human material perception. Notably, many of these dimensions mapped onto shape-related cues, such as long, thin, round, and bulbous. This suggests that, even when judging static images, observers rely on shape to group materials together. 2) Moderate correlations suggest that additional underlying factors influence perceptual judgments beyond the motion and shape statistics we considered in our analysis. Our estimates of motion and shape statistics don’t precisely capture the timings of deformations or interactions with external forces. However, when an observer views these animations, even in dot and line conditions, they can extract precise information about events, including timings and what caused them (Ji & Scholl, 2024). Simple shape and motion statistics differences cannot capture such detailed differences. Thus, to understand material perception fully, we should incorporate interactions with external forces and interactions between shape and motion statistics at each time point. 3) Higher correlation values for line and dot conditions suggest that when information about optical material properties is limited, participants increasingly depend on motion and shape cues to estimate material properties, with the line condition best reflecting the differences in underlying motion and shape statistics.

The primary aim of this analysis was exploratory, to examine how motion and shape statistics vary across different material categories rather than pinpointing exact statistical features used by the visual system. Many observed correlations are moderate, and several statistics can be interrelated. For instance, materials with greater surface area fluctuations often exhibit higher speed as well. Furthermore, our stimulus set included a diverse range of event contexts, e.g., objects falling, a jelly cube being cut, and a fabric flag flowing through the air. Each of these events involves unique interactions that extend beyond material properties alone. While such variability underscores the robustness of the visual system in inferring material properties from dynamic cues, it also complicates the interpretation of motion and shape statistics. Motion or shape statistics are not solely determined by material properties but are also influenced by event context; for example, a rigid object thrown against a wall exhibits motion characteristics shaped by impact force, whereas the same object dragged across a floor follows a completely different motion profile. Yet, despite these variations, the visual system successfully recognizes objects in both scenes. This highlights that motion information is a powerful cue, enabling the visual system to simultaneously identify both the nature of the event and the material involved.

Indeed, previous research has demonstrated that the visual system effectively utilizes motion trajectories to extract information about events (Bingham et al., 1995; McCONNELL et al., 1998; Muchisky & Bingham, 2002; Runeson, 1976; Wickelgren & Bingham, 2001). Runeson (1976) introduced the *Kinematic Specification of Dynamics (KSD)* framework, proposing that observers use kinematic information to infer dynamic properties of events. Building on this, Bingham et al. (1995) isolated motion information using patch-light displays of nine events spanning different dynamic categories, including rigid-body dynamics (e.g., free fall, pendulum motion, rolling ball, struck ball), biodynamics (e.g., hand-moved spring, hand-moved pendulum), hydrodynamics (e.g., stirred water, splash), and aerodynamics (e.g., falling leaves). Their findings showed that observers could successfully recognize each event based purely on motion trajectories. While they did not specifically focus on material perception, it provides valuable context for our research, reinforcing the role of dynamics in both event and material perception. Understanding how these two dimensions interact could offer deeper insights into the mechanisms underlying visual inference of physical properties. A promising direction for future research is to systematically simulate a range of events across different materials and examine how shape and motion interact to give information about the material in each context or event separately.

## Conclusions

We conclude that dynamic line drawings capture material qualities in a range of different material categories. We demonstrated that contour motion, including both occluding contour motion (OCM) and internal contour motion (ICM), plays a critical role in material perception. Motion and shape statistics derived from 2D animations only partially explain the perceptual differences between material categories, highlighting a significant gap to be addressed in future research.

## Acknowledgements

This work was supported by the Deutsche Forschungsgemeinschaft (German Research Foundation, DFG) project number 222641018 – SFB/TRR 135 and DFG under Germany’s Excellence Strategy (EXC 3066/1 “The Adaptive Mind”, Project No. 533717223).

## Supplementary Figures

**Figure S1.**
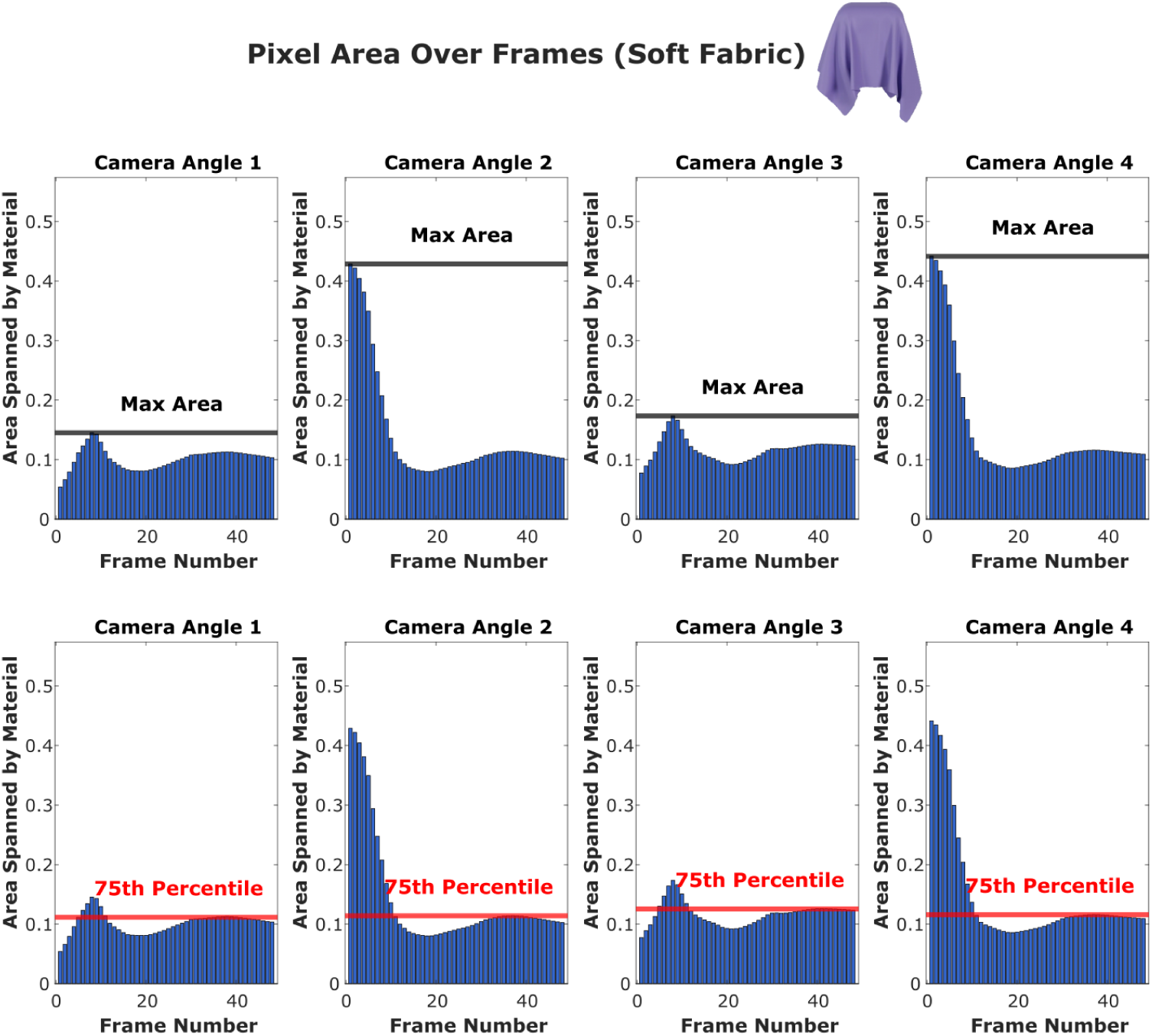
The area spanned by the material across frames for one of the exemplars (soft fabric) viewed from four different camera angles. Each bar represents the proportion of pixels occupied by the material in a given frame. The top row highlights the maximum area (black line), which varies significantly across different camera angles due to differences in viewpoint and the distribution of the area per frame. Some views exhibit a relatively normal distribution of area spanned, while others are heavily skewed, leading to much higher maximum values. The bottom row highlights the 75th percentile area (red line), which remains relatively consistent across views. This choice ensured that the number of sampled vertices remained comparable, avoiding over-representing frames with extreme values and maintaining a more uniform dot density across different animations.

## Appendix A: Rendering Details

This appendix provides rendering details for each exemplar across all categories in Blender.

***1. Rigid-breakable:*** This category included four exemplars depicting hard objects composed of materials that shattered upon impact with a rigid surface. The representative frames are shown on the bottom left panel of Figure 3. Two exemplars were rigid objects thrown against an invisible wall, while the other two were rigid objects dropped from an elevation onto an invisible ground. We used the Cell Fracture add-on in Blender to shatter objects into pieces, and rigid body physics to simulate their realistic interactions upon impact. The objects thrown at the wall were identical, oval-shaped, grey objects. The impact with the wall occurred in the 7th frame in both animations, causing the object to break into 7 or 8 pieces. The difference between the two animations was that the object was rendered as hollow in one and as solid-filled in the other. The objects dropped on the ground were cubes. The impact occurred in the 12th frame in both animations, breaking the cube into 7 or 8 pieces upon impact. Consistent with the prior set, one animation consisted of the hollow cube, and the other consisted of the solid-filled cube.
***2. Smoke:*** The stimulus set consisted of three animations (top right panel of Figure 3). Two animations show smoke: one with smoke spreading uniformly from the center, while the other displays a more dynamic dispersal pattern starting from the center. The third animation shows mist that emits from a defined source and is dispersed within an invisible box-like domain. We used Blender’s particle system in combination with fluid simulation, simulating the movement of particles through a fluid field.
***3. Liquid:*** The stimuli for the liquid category consisted of three exemplars: two showing liquids of different viscosities constrained within a box-like domain, where the perturbed liquid flows back and forth within the boundaries of the container, and one featuring liquid droplets falling freely and splashing upon impact with the ground. The representative frames are shown in the top middle panel of Figure 3. These simulations were created using Blender’s fluid and particle systems.
***4. Jelly:*** This category consisted of four exemplars featuring objects made of jelly, as shown in the top left panel of Figure 3. Two animations contained cube-shaped objects: one was dropped onto an invisible ground, and the other was sliced in half by an invisible knife. The third animation depicted a jelly sheet that collided with an invisible cube object when dropped onto it, bouncing up and down. The final exemplar was a potato-shaped jelly object that was also dropped onto an invisible ground. We used Blender’s particle system and the Molecular add-on to simulate jelly dynamics.
***5. Fabric:*** This category included four animations, as shown in the bottom right panel of Figure 3. These included a soft cloth that draped over an invisible cube when dropped, a hard cloth that similarly fell over an invisible cube but retained a more rigid structure and bounced, a flag that fluttered in the wind, and a soft fabric ball filled with air that is dropped onto an invisible ground, deforming upon impact. Fabric dynamics were simulated using Blender’s cloth simulation system.

## Appendix B: Calibration procedure

The calibration procedure was based on the virtual chinrest method developed by Li et al. (2020). To measure screen size, participants completed a “card task,” which involved adjusting an on-screen image of a credit card to match the physical size of an actual card placed on the screen. This adjustment was done using a slider, and the resulting card width in pixels was recorded. The procedure was repeated three times, and the average width was used to calculate the Logical Pixel Density (LPD), defined as pixels per millimeter. LPD was then used to scale the animations, ensuring a consistent physical size across different displays.

Participants were instructed to sit at an approximate viewing distance of 50 cm. However, to measure the actual distance being set, participants completed a *‘blind spot task’*. In this task, a stationary black square was presented on the right side of the screen while a red dot moved horizontally from right to left. Participants were required to cover their right eye and fixate on the stationary black square. They were instructed to press the spacebar when they perceived the dot disappearing from their sight. The distance between the center of the black square and the red dot is recorded at the moment the spacebar is pressed. The procedure was repeated 5 times, and the average of this distance was taken as the final measurement, which was then used to estimate viewing distance. Please refer to Li et al. (2020) for further details of the calibration procedure and measurements.

## Appendix C: Construction of RDMs from the similarity judgment task

In a triplet 2-AFC similarity judgment task, material similarity is defined as the probability *P*(*i*, *j*) of participants choosing material animations (*i*) and (*j*) as belonging together, marginalized across all contexts imposed by the third animation (*k*). To generalize across all contexts, we needed to test each unique combination of exemplars in a triplet setting. For that, we took each of the 18 exemplars from the set of animations as a reference and paired them with every possible unique combination of exemplars from material categories distinct from the reference category. Triplets containing exemplars from the same category were excluded, as we were only interested in cross-category similarity judgments. This led to a total of 1,380 unique triplets. The whole set is then divided into four subsets with 345 triplets in one subset, which were presented in four separate runs. The order of the triplet presentations was randomized across trials within each run. For every trial, the three animations in a triplet were presented from a randomly selected camera angle, chosen from six to eight pre-rendered options. Each run consisted of four blocks, taking an average of 40 minutes to complete. Trial numbers and block numbers were displayed on the bottom-right and bottom-left corners of the screen, respectively, for participants’ reference.

Four runs were required to complete the entire set of 1380 unique triplets. Since it could be very tedious to complete for one participant, we provided participants with the option to choose the number of subsets they would prefer to take part in. Eighty participants completed all four runs; 3 participants completed three runs; 14 participants completed two runs; and 3 participants completed only a single run. Importantly, if a participant completed multiple runs, all were presented in the same rendering condition. Consequently, we achieved 30 repetitions for each complete set of triplets associated with each rendering condition: full, line, and line.

To estimate the perceived similarity between materials, we calculated the frequency with which a reference exemplar (*i*) was paired with each of the test exemplars (*j*) across all unique triplet combinations. This count was normalized by dividing it by the total number of unique combinations for each reference exemplar, resulting in the probability of grouping *i* and *j* together (*P*(*i*, *j*)), generalized across all contexts (*k*). To account for symmetry, we combined *P*(*i*, *j*) and *P*(*j*, *i*) yielding a unified probability of pairing materials *i* and *j* that ranged from 0 to 2. Finally, we subtracted these values from 2 to generate a dissimilarity matrix, which provided the required RDMs (Representational Dissimilarity Matrices) showing perceived dissimilarity between materials.

## Notes

### Competing Interest Statement

The authors have declared no competing interest.

### Summary of Updates

Author affiliations are updated in the current version.

